# The *SCN1A* Philadelphia variant – a gain-of-function mutation causing an early-onset epileptic encephalopathy

**DOI:** 10.1101/2022.06.29.498154

**Authors:** Jérôme Clatot, Shridhar Parthasarathy, Stacey Cohen, Jillian McKee, Shavonne Massey, Ala Somarowthu, Ethan M. Goldberg, Ingo Helbig

## Abstract

**Objective:** Loss-of-function variants in *SCN1A* cause Dravet Syndrome, the most common genetic developmental and epileptic encephalopathy (DEE). However, emerging evidence suggests separate entities of *SCN1A*-related disorders due to gain-of-function variants. Here, we aim to refine the clinical, genetic, and functional electrophysiological features of a recurrent p.R1636Q gain-of-function variant, identified in four individuals at a single center.

**Methods:** Individuals carrying the recurrent *SCN1A* p.R1636Q variant were identified through diagnostic testing. Whole-cell voltage-clamp electrophysiological recording in HEK-293T cells was performed to compare the properties of sodium channels containing wild-type Nav1.1 or Nav1.1-R1636Q along with both Navβ1 and Navβ2 subunits, including response to oxcarbazepine. To delineate differences to other *SCN1A*-related epilepsies, we analyzed electronic medical records.

**Results:** All four individuals had an early-onset DEE characterized by focal tonic seizures and additional seizure types starting in the first few weeks of life. Electrophysiological analysis showed a mixed gain-of-function effect with normal current density, a leftward (hyperpolarized) shift of steady-state inactivation, and slower inactivation kinetics leading to a prominent late sodium current (*I*_*Na*_). The observed functional changes closely paralleled effects of pathogenic variants in *SCN3A* and *SCN8A* at corresponding positions. Both wildtype and variant exhibited sensitivity to block by oxcarbazepine, partially correcting electrophysiological abnormalities of the *SCN1A* p.R1636Q variant. Clinically, a single individual responded to treatment with oxcarbazepine. Across 51 individuals with *SCN1A*-related epilepsies, those with the recurrent p.R1636Q variants had the earliest ages of onset.

**Interpretation:** The recurrent *SCN1A* p.R1636Q variant causes a clinical entity with a wider clinical spectrum than previously reported, characterized by ultra early-onset epilepsy and absence of prominent movement disorder. Functional consequences of this variant lead to mixed loss- and gain-of-function that is partially corrected by oxcarbazepine. The recurrent p.R1636Q variant represents one of the most common causes of early-onset *SCN1A*-related epilepsies with separate treatment and prognosis implications.

**Key Points:**

1. Loss-of-function variants in *SCN1A* cause Dravet syndrome, but gain-of-function variants have an emerging clinical spectrum.
2. The *SCN1A* p.R1636Q variant shows similar overall gain-of-function effects to identical missense variants in other voltage-gated sodium channels.
3. Features of four unreported individuals with *SCN1A* p.R1636Q from a single center expand the *SCN1A* gain-of-function phenotype.
4. Individuals with this variant are recognizable by their ultra early-onset seizures in contrast to Dravet syndrome.

## Introduction

The developmental and epileptic encephalopathies (DEE) are severe epilepsies with a range of comorbidities.^1,2^ Most DEE start in early infancy and are typically associated with developmental delays and intractable seizures.^3,4^ More than 100 genetic causes have been identified in the last two decades, with *SCN1A, STXBP1, SCN2A*, and *KCNQ2* as some of the most common genes.^5-8^ Among the epilepsy genes, *SCN1A* is the most common genetic cause for DEE with an expected frequency or 1:20,000 or more.^9^

Loss-of-function variants in *SCN1A* are known to cause a range of fever-associated epilepsies including Dravet Syndrome and Genetic Epilepsy with Febrile Seizures Plus (GEFS+).^9,10^ A molecular diagnosis of a loss-of-function variant in *SCN1A* is already valuable to guide clinical decision-making. For example, sodium channel blockers such as oxcarbazepine, phenytoin, and lamotrigine, are strictly contraindicated in Dravet Syndrome early in the disease course, as they may cause seizure exacerbation.^11^ In contrast, ASMs such valproic acid, clobazam, stiripentol, cannabidiol, and fenfluramine are effective.^11-14^ Additionally, loss-of-function variants in *SCN1A* are promising targets for precision medicine efforts including anti-sense oligonucleotide (ASO) therapy.^15,16^

In contrast to the well-established loss-of-function phenotypic spectrum, *SCN1A*-related epilepsies due to gain-of-function variants represent a nebulous clinical entity. Individuals with such variants have been reported to have clinical features distinct from Dravet Syndrome, including familial hemiplegic migraine, hyperkinetic movement disorders, and an earlier-onset epileptic encephalopathy. A recent study highlighted the range of gain-of-function *SCN1A*-related disorders, including the recurrent *SCN1A* p.R1636Q variant in 5 individuals recruited across a large collaborative network, suggesting a prominent movement disorder phenotype as a common feature in addition to severe intellectual disability.^17^

Here, we describe four previously unreported individuals with the recurrent *SCN1A* p.R1636Q variant identified through routine care. All individuals had seizures starting in the first weeks of life; seizure types and onset clearly stands out from other *SCN1A*-related epilepsies. Electrophysiological properties of this variant reveal an overall gain-of-function effect, with underlying mixed gain- and loss-of-function changes, as the disease mechanism. All four individuals were identified at a single healthcare network within less than two years. Therefore, we refer to *SCN1A* p.R1636Q as the “Philadelphia variant” to raise awareness for the emerging role of *SCN1A* gain-of-function variants, which may require therapeutic approaches opposed to those for Dravet Syndrome.

## Materials & Methods

### Genetic analysis and clinical assessment

Individuals with the recurrent *SCN1A* p.R1636Q variant were identified and assessed through routine clinical care within the Epilepsy NeuroGenetics Initiative (ENGIN) at Children’s Hospital of Philadelphia. Genetic testing was performed clinically through gene panel testing and/or trio-based whole exome sequencing. Clinical data for all individuals was accessed and reviewed using medical records, including neuroimaging and EEG data. Informed consent for participation in this study was obtained from subjects themselves or parents of all probands in agreement with the Declaration of Helsinki, and the study was completed per protocol with local approval by the Children’s Hospital of Philadelphia (CHOP) Institutional Review Board (IRB 15-12226).

### Molecular Biology and plasmid constructions

Wild-type (WT) and variant human *SCN1A* cDNA (isoform 1; reference sequence NM_001165963.4) were synthetized and subcloned in pcDNA3.1, resulting in a pcDNA3.1-*SCN1A* plasmid. *SCN1B* and *SCN2B* genes cDNA (NM_001037.5 and NM_004588.5) encoding the human Na+ channel accessory subunits Na_v_β1 and Na_v_b2 were subcloned in a pCMV-FusionRED plasmid resulting in a tricistronic pCMV-FusionRED/*SCN1B*/*SCN2B* plasmid.

### Cell culture and transfection

Cell culture, transfections, and electrophysiological experiments were performed using HEK-293T cells. Cells were grown at 37°C with 5% CO_2_ in Dulbecco’s modified Eagle’s medium (DMEM) supplemented with 10% fetal bovine serum, 2 mM L-glutamine, and penicillin (50 U/mL)-streptomycin (50 μg/mL). HEK-293T cells were co-transfected with 0.15 µg of pCMV-Fusionred/*SCN1B*/*SCN2B* and 2 µg pcDNA3.1-*SCN1A* WT using 10 uL of PolyFect transfection reagent (QIAGEN; Germantown, MD, U.S.A.) in 35-mm culture dishes following the manufacturer’s instruction. Cells were then incubated for 48 hours after transfection prior to electrophysiological recording. Transfected cells were then trypsinized and seeded on 35mm dishes to a density allowing single red positive cells to be identified. After 4 hours at 37 °C in 5% CO_2_ cells were washed once with extracellular Tyrode’s solution for 5 minutes prior recording.

### Voltage-clamp electrophysiology

Patch-clamp recordings were carried out in the whole-cell configuration at room temperature (*∼*22°C). Cells were bathed in an extracellular Tyrode’s solution containing (in mM): 150 NaCl, 2 KCL, 1 MgCl2, 1.5 CaCl2, 1 NaH2PO4, 10 glucose, 10 HEPES, adjusted to pH 7.4 with CsOH. Intracellular solution was (in mM): 35 NaCl, 105 CsF, 2 MgCl2, 10 EGTA, 10 HEPES, adjusted to pH 7.4 with CsOH. For experiment requiring the perfusion of TTX and/or oxcarbazepine (Tocris Bioscience), a Biopen nanoperfusion system was used allowing the perfusion of a single cell. Stock solutions of TTX (2 µM in water) and oxcarbazepine (25 mM in 100% DMSO) were first prepared, and compounds were then diluted in their final concentration in extracellular Tyrode’s solution. Control experiments were performed using Tyrode’s solution with the corresponding vehicle without added compound.

Ionic currents were recorded with an Axopatch 200B amplifier (Axon Instruments, CA, USA). Patch pipettes (Corning Kovar Sealing code 7052, WPI) had resistances of 1.5-2.5 MΩ when filled with intracellular solution. Currents were filtered at 10 kHz (−3 dB, 8-pole low-pass Bessel filter) and digitized at 50 kHz (NI PCI-6251, National Instruments, Austin, TX, USA). Data were acquired using Clampex and analyzed with Clampfit 11.1 (Molecular Devices, CA, USA) and MATLAB (MathWorks).

Current-voltage relationships (I/V curves) were constructed by eliciting test potentials to -80 mV to 55 mV in increments of 5 mV from a holding potential of -90 mV at 0.2 Hz frequency. The steady-state inactivation protocol was established from a holding potential of -120 mV and a 500-ms conditioning prepulse was applied in 5 mV increments from -140 and 5 mV, followed by a 400-ms test pulse to 0 mV. Voltage-dependence of activation and steady-state inactivation curves were fitted to the Boltzmann equation:

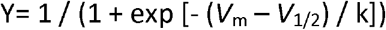

where *V*_*m*_ is the membrane potential, *V*_*1/2*_ is the voltage of half-maximal activation or channel availability, and k is the inverse slope factor. For activation curves, Y represents the relative conductance and k is > 0; for availability, Y represents the relative current (*I*_*Na*_/*I*_*Na*_max) and k is < 0. Inactivation kinetics time constants (*τ*_fast_ and *τ*_slow_) were calculated by fitting with a double exponential function using Clampfit. Persistent current (measured via subtraction after application of TTX) and sensitivity to oxcarbazepine were measured using a train depolarization pulses to 0mV for 500ms ms from a holding potential at -120 mV at a frequency of 0.6 Hz.

### Electronic Medical Record (EMR) analysis

Documented seizure onset was retrieved for all four individuals with the recurrent *SCN1A* p.R1636Q and 48 additional individuals with *SCN1A*-related epilepsies enrolled in the Epilepsy Genetics Research Protocol (EGRP, IRB 15-12226) from our study database. Electronic medical records (EMR) were also retrieved for all individuals within our healthcare network with a mention of “*SCN1A*” in full text patient notes and a clinical diagnosis of epilepsy, defined by at least one patient encounter coded with ICD10 G40, R56, P90 or ICD9 345, 780.3, 779, the most common epilepsy codes. We referred to this combination of billing codes as G40+. All patient encounters were extracted and time-stamped to assess time course and completeness of the medical records. Analysis and visualization of clinical and EMR data was performed through R Studio (R Foundation for Statistical Computing, Vienna, Austria).^18^ Data extraction and analysis was performed within the HIPAA-compliant Arcus framework at Children’s Hospital of Philadelphia. Arcus is an initiative of the Research Institute to link clinical and research data to accelerate science and was used for data analysis as previously described.^19^

### Statistical analysis

Data are presented as mean ± SEM. Statistical significance was estimated with SigmaPlot® software using one-way ANOVA, as appropriate; significance was set at *p* < 0.05. All recordings and analysis were performed blind to identity of the variant via assignment of a randomization code.

## Results

### *SCN1A* p.R1636Q results in a DEE different from Dravet Syndrome

We identified four individuals through clinical care with the recurrent *SCN1A* p.R1636Q variant. All individuals identified clinically within our study had a common clinical presentation with early onset epilepsy, typically starting in the neonatal period. We also reviewed the clinical features of eight previously reported individuals with the recurrent *SCN1A* p.R1636Q variant in the literature (Table 1).^17,20-23^

**Table 1.** Clinical features in three individuals with the recurrent variant p.R1636Q of Na_V_1.1. A: Atonic, Ab: Absence, At A: atypical absence, DD: developmental delay, EPC: epilepsia partialis continua, ESES: electrical status epilepticus in sleep, F: focal, FD: focal disharges, FS: febrile seizures, GSW: Generalized spike-wave, HC: hemiclonic, ID: intellectual disability, IS: infantile spasms, LTG: lamotrigine, M: myoclonic, MF: multifocal, NA: non-ambulatory, NCSE: non-convulsive status epilepticus, NV: non-verbal, OXC: oxcarbazepine, PHT: phenytoin, PSW: polyspike wave, SE: status epilepticus, T: tonic, TC: tonic-clonic, UL: upper limb *: deceased

|  | Individual 1 | Individual 2 | Individual 3 | Individual 4 | Individual 5 | Individual 6 | Individual 7 | Individual 8 | Individual 9 | Individual 10 | Individual 11 | Individual 12 |
| --- | --- | --- | --- | --- | --- | --- | --- | --- | --- | --- | --- | --- |
| Source | Butler (2017) | Nair (2018) | Alame (2019) | Harkin (2007); Brunklaus (2022) | Brunklaus (2022) | Brunklaus (2022) | Brunklaus (2022) | Brunklaus (2022) | This study | This study | This study | This study |
| Age (sex) | N/A | N/A | N/A | 20y* (male) | 8y* (female) | 11y (male) | 23y (female) | 5y (male) | 13m (male) | 4y (male) | 19y (female) | 35m (male) |
| Variant | c.4907G>A, p.R1636Q unknown | c.4907G>A, p.R1636Q unknown | c.4097G>A, p.R1636Q *de novo* | c.4907G>A, p.R1636Q *de novo* | c.4907G>A, p.R1636Q unknown | c.4907G>A, p.R1636Q *de novo* | c.4907G>A, p.R1636Q *de novo* | c.4907G>A, p.R1636Q *de novo* | c.4907G>A, p.R1636Q *de novo* | c.4907G>A, p.R1636Q *de novo* | c.4907G>A, p.R1636Q, not paternally inherited | c.4907G>A, p.R1636Q *de novo* |
| Seizure types (onset of first seizure) | M, otherwise unknown | “early seizures”<br>“Dravet syndrome” | “Dravet syndrome” | GTC (3w), Ab, AtA, T, F, IS, M, HC | T (3m), F, GTC | TC (2m), C, F, T, M, SE | TC (10w), M, HC, T, SE, At | TC (2m), F, SE, T, HC | T (2w) | GTC (DOL3), T (infancy) | T (infancy) | TC (neonatally), F (2 months), SE (4 months), A (12 months) |
| EEG | N/A | N/A | N/A | MF discharges, FD, ESES | GSW, PSW, NCSE | GSW, FD, EPC | MF discharges, ESES, NCSE | MF discharges, FD | FD | MF discharges, ESES | N/A | MF spikes |
| Brain MRI | N/A | N/A | N/A | Normal 5y and 18y | Normal 5m, mild atrophy 1.5y | Normal 2y and 5y | Normal 3y and 6y | Normal 1.5y | Normal 8w | Normal 1m and 1.5y | Normal 2.5y and 4y | Normal 4m |
| Neurological/physical exam | Dystonia, spasticity | N/A | N/A | Hypotonia, right UL weakness & posturing (18 m), ataxic (2.5 y), crouch gait (6 y), wheelchair bound (20 y) | Hypotonia, chorea of UL, myoclonus (1 y), Profound ID, NV, NA (2.5y → regression), acquired microcephaly, bone fractures, eyelid myoclonia | Dystonia, dyskinesia (7y), delay (1y) and regression (7y), profound ID, NA, NV, acquired microcephaly, eyelid myoclonia | Dyskinesia of right UL (2y), delay (1-2y) and regression (5y), severe ID, hypotonia | Head tremor (2y), intermittent ataxia, delay (1-2y), severe ID, single words, ambulant, autism-like features | Normal | Axial hypotonia, NA, NV, bilateral chorea, profound DD | NV, broad-based gait, severe ID | Axial hypotonia, brisk reflexes unilaterally (left) |
| Sodium channel blocker AEDs (effect) | N/A | N/A | N/A | LTG (seizure reduction), PHT (seizure reduction) | PHT (seizure reduction) | PHT (no effect) | LTG (seizure worsening), PHT (seizure reduction) | OXC (seizure freedom, temporary) | OXC (seizure freedom) | None | None | None |

Individual 1 was a 14-month old male with focal epilepsy. He presented with focal tonic seizures on day-of-life 14 which resolved with oxcarbazepine monotherapy with a single episode of breakthrough seizures with preceding vaccinations (Figure 1). Treatment with oxcarbazepine had been initiated prior to genetic testing and was maintained after the genetic diagnosis. At the last clinical assessment at 14 months, neurological exam was unremarkable, and all developmental milestones were within the typical range. EEG performed at 6 weeks was unremarkable, but EEG at 8 weeks showed focal interictal discharges and one focal seizure. MRI Brain performed at the age of 8 weeks was unremarkable. Gene panel testing at 4 months identified the SCN1A p.R1636Q variant which was confirmed *de novo*.

**Figure 1.**
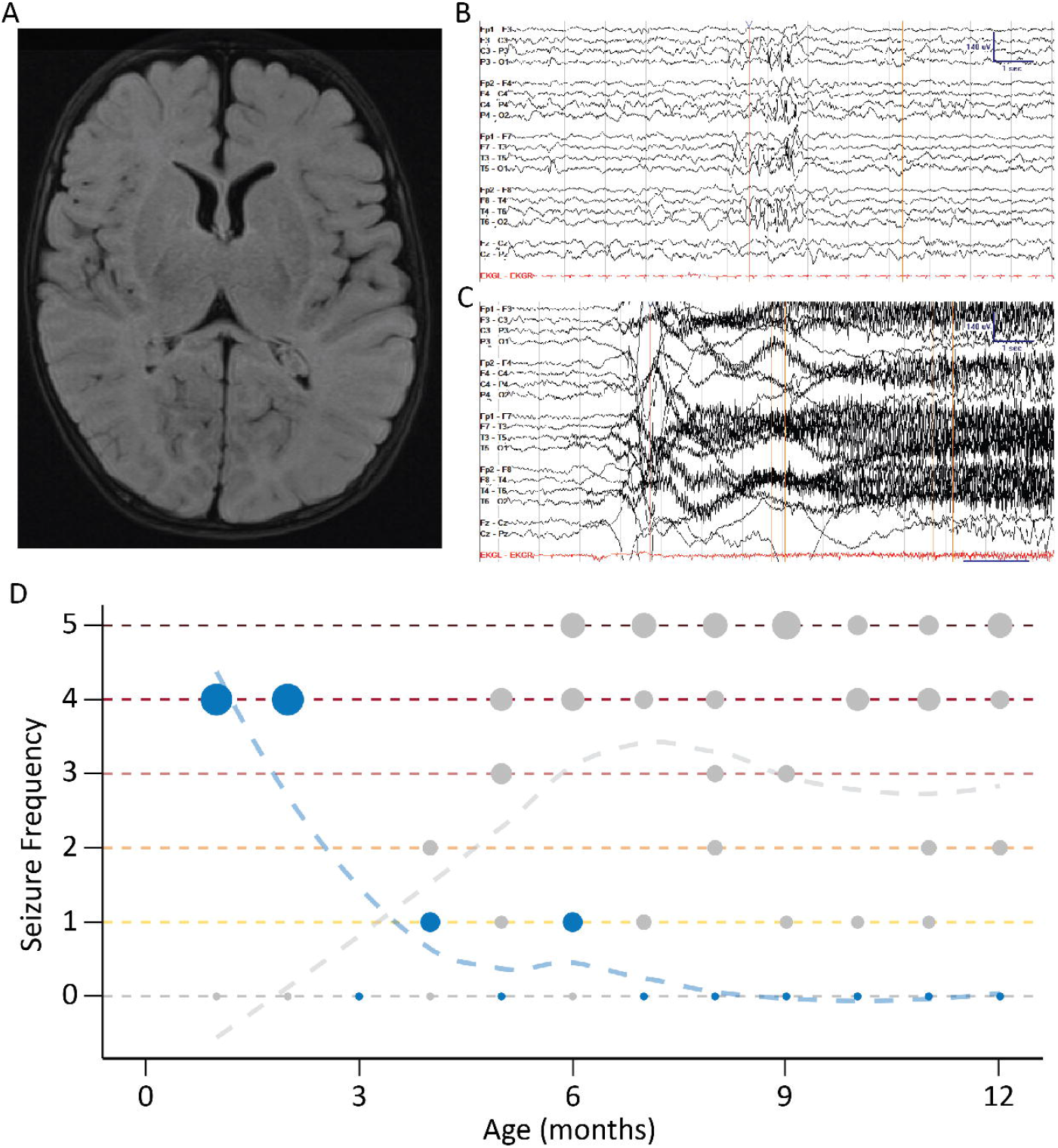
Representative clinical features for one individual with *SCN1A* p.R1636Q, which is different from Dravet Syndrome. **(A)** Axial T2 FLAIR MRI acquired at 2 months of age. **(B)** Independent P3/P4 sharps, recoded in bipolar montage and sensitivity of 7 uV/mm. **(C)** Onset of tonic seizure, with fast activity originating at P4/Cz and spreading bilaterally. This is associated clinically with a cry then a tonic followed by clonic phase lasting 1.5 minutes. **(D)** Seizure history across first year of life compared to five demographically matched individuals with Dravet Syndrome using standardized seizure frequency labels of the Pediatric Epilepsy Learning Health System: 0 = seizure free, 1 = monthly, 2 = weekly, 3 = daily, 4 = many per day, 5 = several per day. Seizure onset in this individual was earlier than would be expected in Dravet syndrome.

Individual 2 was a 3-year-old male with early onset intractable epilepsy and profound global developmental delay. He initially presented with bilateral tonic-clonic seizures on day-of-life 3, which required hospital admission and treatment with phenobarbital. He continued to have bilateral tonic-clonic seizures until age 2 years. During infancy, he developed tonic seizures. At the last clinical assessment at 33 months, he had been seizure free for four months and was treated with a combination of cannabidiol, levetiracetam, topiramate, and clobazam. In addition to seizures, Individual 2 had prominent dyskinesia and choreoathetosis, which was first noted at 3 years. The development was unremarkable in early infancy apart from hypotonia. At the age of 2.5 years, a notable developmental regression was observed, which coincided with an increase in seizures. At the last clinical assessment at 33 months, the neurological exam demonstrated severe axial and truncal hypotonia and bilateral chorea. He had profound developmental delay and was non-ambulatory and nonverbal. EEG monitoring was first performed on day-of-life 2 days and was unremarkable. By 4 months, multifocal epileptiform activity was captured, but a follow-up EEG was 8 months was unremarkable. Long-term EEG monitoring at 16 months again showed multifocal epileptiform discharges that activated in sleep with diffuse background slowing. On the most recent EEG at the age of 2.5 years, diffuse background slowing and electrical status epilepticus in sleep (ESES) that occupied 85% of non-REM sleep was noted. MRI Brain performed at the age of 2.5 years was unremarkable. Trio-based whole exome sequencing at the age of 2.5 years identified the *SCN1A* p.R1636Q variant which was confirmed *de novo*.

Individual 3 was a 19-year old woman with chronic static encephalopathy and intractable epilepsy. Tonic seizures allegedly started in early infancy, but early medical records were not available for review. She presented with recurrent events of abnormal breathing and cyanosis shortly after birth, which were identified as seizures on EEG. Seizures were intractable to treatment with oxcarbazepine, levetiracetam, gabapentin, and topiramate but frequency was significantly reduced with valproic acid. She continued to have tonic seizures throughout childhood and adolescence, which typically occurred out of sleep. Developmental milestones were delayed globally since early infancy. She started walking at 3-4 years with a broad-based gait. She also received a diagnosis of autism spectrum disorder in early childhood. Spoken language was never acquired. She had severe intellectual disability, and episodes of regression were not observed. At the last clinical assessment at 18 years, she had infrequent nocturnal tonic seizures and was treated with valproic acid. The neurological exam was characterized by moderate hypotonia, but not focal neurological signs. EEG monitoring in infancy confirmed focal seizures, and further EEGs were not performed by family preference. Brain MRI performed at 31 months and 47 months were unremarkable. Gene panel testing at the age of 18 years identified the *SCN1A* p.R1636Q variant, which was not paternally inherited.

Individual 4 was a 35-month-old boy with global developmental delay, intractable epilepsy, and submucosal cleft palate. He had episodes starting at day of life 1 characterized by bilateral stiffening, foaming, and turning red. Initial EEG recordings were unremarkable and he was treated with antacids under assumption of gastroesophageal reflux. The episodes continued to occur daily and a repeat EEG at the of 4 months showed six seizures arising from bi-posterior/parasagittal and right posterior temporal regions, confirming the diagnosis of epilepsy. In retrospect, given the unchanged presentation of these events until EEG confirmation at 4 months, it is assumed that seizures started neonatally. In addition to the tonic-clonic seizures with presumed neonatal onset, he also had episodes with behavioral arrest, lip smacking, and pupillary dilation starting at 2 months and occurring daily since. These episodes were not captured by EEG, but were considered focal impaired awareness seizures upon review of video recordings by the treating physician. He had two episodes of status epilepticus, one in the setting of aspiration pneumonia at 4 months and one in the setting of medication wean at 33 months. He walked at 18 months and spoke his first word at 2 years, with 30 single words and no phrases obtained by age of last evaluation, 35 months. At the last visit at 35 months, the neurological exam was characterized by axial hypotonia and brisk deep tendon reflexes on the left side. Most recent EEG at 29 months showed rare spikes in the left occipital region and bilaterally in the central region. MRI Brain performed at 4 months was unremarkable. Gene panel testing at the age of 4.5 months identified the *SCN1A* p.R1636Q, which was confirmed *de novo*.

In contrast to the eight individuals previously reported with the recurrent SCN1A p.R1636Q variant, the clinical presentation of our patients was milder. Only one individual had a hyperkinetic movement disorder and developmental delay was mild in 2/4 individuals.

### Electrophysiological analysis of *SCN1A* p.R1636Q shows overall gain-of-function

To investigate the disease mechanism of *SCN1A* p.R1636Q, we expressed the variant in HEK-293T cells for characterization via whole-cell voltage clamp recording. We recorded wild-type or variant h*SCN1A* cDNA was co-expressed with wild-type β1 and β2.

We first measured peak current density and established the current-voltage relationship (I-V curve). We found no significant difference in peak current density between cells expressing *SCN1A-*WT and p.R1636Q, being -90.0 ± 16.3 pA/pF for WT (*n* = 15) and -126.9 ± 24.5 pA/pF for p.R1636Q (*n* = 12; *p* = 0.407 via one-way ANOVA), respectively, at -5 mV (**Figure 2A-B**). Furthermore, we observed an approximate -5 mV (hyperpolarizing) shift of the voltage-dependence of activation, from -21.2 ± 1.4 mV for Nav1.1-WT to -27.2 ± 1.4 mV for Nav1.1-R1636Q (*n* = 16; *p* = 0.009 vs. WT via one-way ANOVA) (**Figure 2C-D)**.

**Figure 2.**
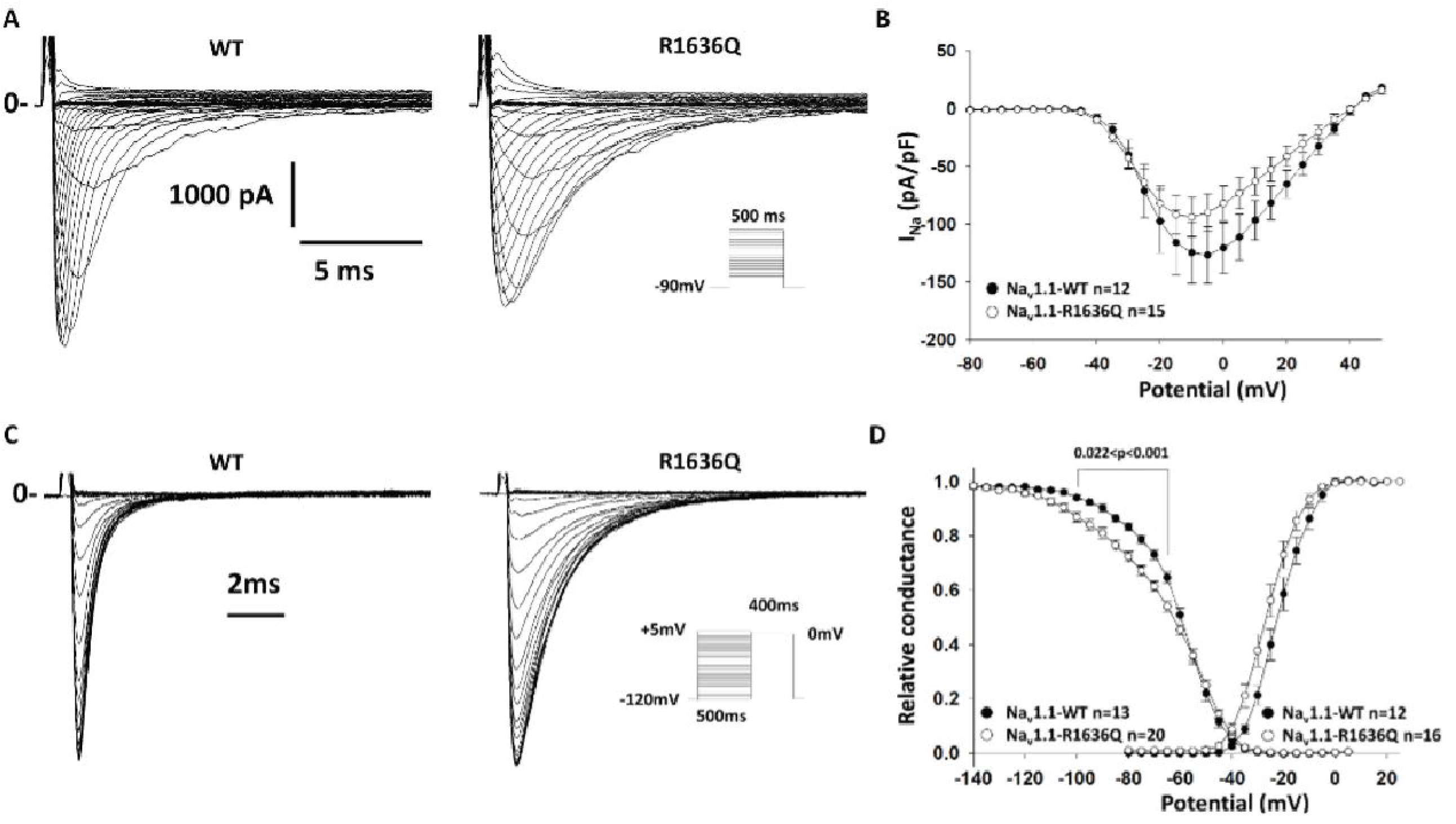
Electrophysiological characterization of Nav1.1-R1636Q reveals normal current density and impaired voltage dependence of inactivation. HEK-293T cells were transfected with Nav1.1-WT or R1636Q along with Navβ1 and Navβ2 subunits. **(A)** Representative Na^+^ current traces for WT or R1636Q in response to voltage steps in the I-V protocol. Solid lines (marked with “0”) indicate the zero current level. **(B)** Averaged current density-voltage relationships (I-V curves). The small difference in mean peak current density was not significant between groups (*p* = 0.407). **(C)** Representative Na^+^ current traces elicited in response to the steady-state inactivation protocol (*inset*). **(D)** Summary data showing normalized voltage dependence of activation and Na+ channel availability/steady-state inactivation. **Note:** The R1636Q variant displays a left-shifted of both voltage-dependence of activation and steady-state inactivation with a significantly decrease slope of voltage-dependence of inactivation.

We then determined the effect of the pR1636Q variant on steady-state inactivation of Nav1.1 channels. We show that Nav1.1-R1636Q displayed a -5 mV hyperpolarized shift of the steady-state inactivation of -66.3 ± 1.1 mV (*n* = 20) compared to WT values of -61.3 ± 1.0 mV (*n* = 13; *p* = 0.004 vs. Nav1.1-R1636Q via one-way ANOVA). The slope k of the steady-state inactivation curve was significantly reduced for the Nav1.1-R1636Q variant, at -3.36 ± 0.20 (*n* = 20) compared to -4.70 ± 0.20 for the WT (*n* = 13; *p <* 0.001 vs. Nav1.1-R1636Q). There was no difference between Nav1.1-WT and Nav1.1-R1636Q in properties of recovery from inactivation.

We next investigated the inactivation kinetics and found that both t_fast_ and t_slow_ of the inactivation kinetics were significantly altered for the variant compared to WT (**Figure 3A-B**). t_fast_ was 2.43 ± 0.21 ms for Nav1.1-R1636Q (*n* = 31) vs. 0.60 ± 0.04 ms for WT (*n* = 20; p < 0.001 vs. Nav1.1-R1636Q); t_slow_ was 15.7± 1.80 for Nav1.1-R1636Q (*n* = 26) vs. 9.36 ± 1.80 for WT (*n* = 19; *p* = 0.003 vs. Nav1.1-R1636Q).

**Figure 3.**
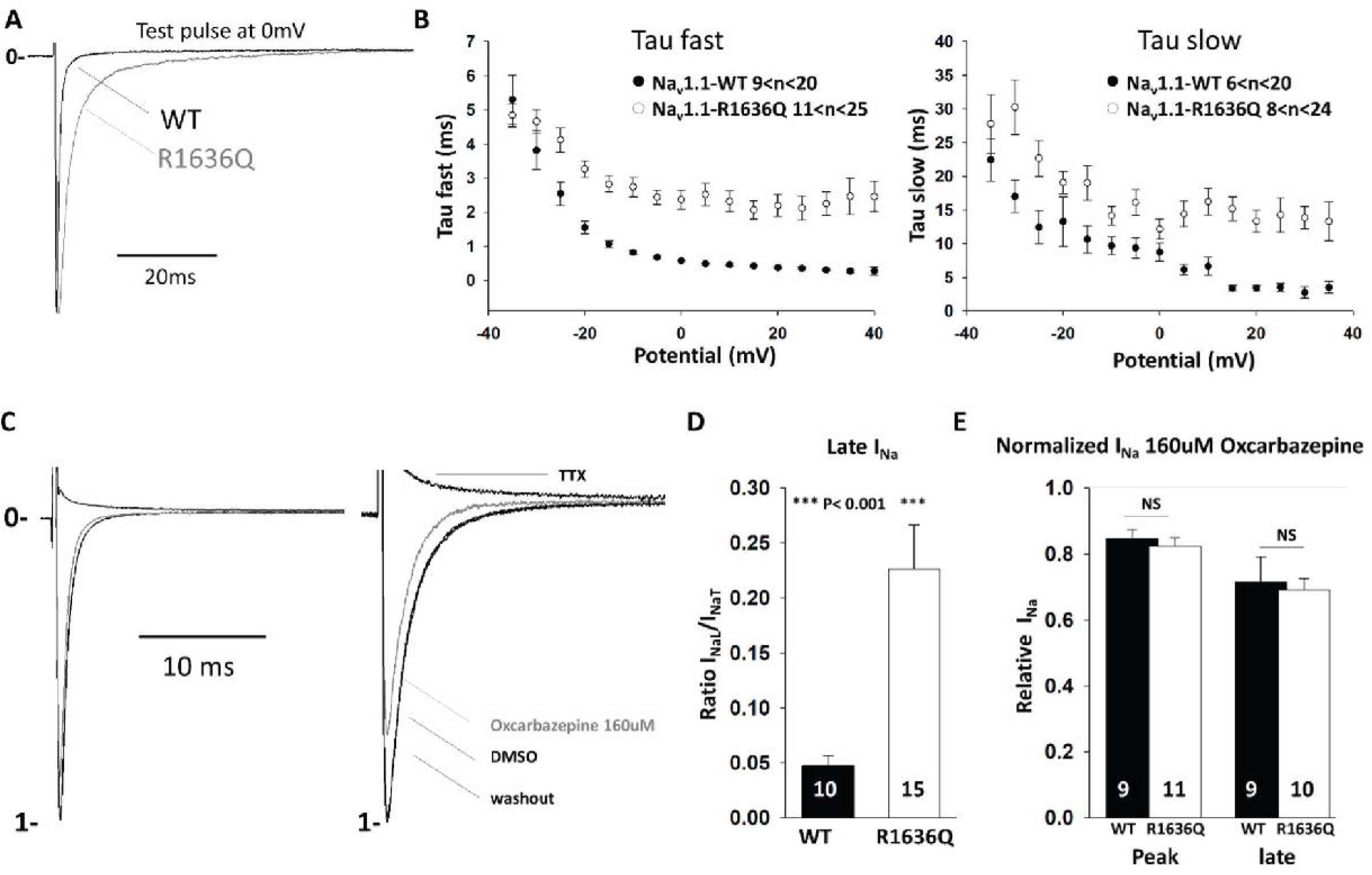
Impaired inactivation and increased persistent current of Nav1.1-R1636Q. HEK-293T cells were transfected with Na_v_1.1-WT or R1636Q along with Navβ1 and Navβ2 subunits. **(A)** Representative I_Na_ traces recorded from cells expressing Nav1.1-WT (*black*) or Nav1.1-R1636Q (*gray*) channels. **(B)** Averaged tau fast and slow values as a function of membrane potential **(C)** Representative *I*_Na_ before and after oxcarbazepine or TTX treatment **(D)** Quantification of *I*_Na_ late (*I*_NaL_) by normalizing INa transient (*I*_NaT_) **(E)** Quantification of remaining *I*_Na_ after treatment with oxcarbazepine (160 µM) at 0 mV from a holding potential of -120 mV.

We then assessed the effect of the Nav1.1-R1636Q variant on persistent Na+ current. We used TTX to quantify the TTX-sensitive slowly-inactivating/persistent Na+ current I_NaP_/I_NaT_ (Figure 3C-D) which revealed a persistent Na+ current of 4.7% ± 0.9 for Nav1.1-WT (*n* = 10) vs. 22.6 ± 4.0% for Nav1.1-R1636Q (*n* = 15; *p* < 0.001).

Based on clinical observation of the treatment response of Individual 1 to oxcarbazepine, we hypothesized that inhibition of persistent current could be achieved using oxcarbazepine, as it is a sodium channel blocker. Indeed, we found that oxcarbazepine efficiently led to significant reduction of peak and persistent I_Na_ of both WT and R1636Q mutant (**Figure 3D-E**). The observed effect of oxcarbazepine on the channel represents partial correction of the gain of function and may explain the seizure reduction achieved in Individual 1 using this medication.

### *SCN1A* p.R1636Q is similar to identical, known disease variants in *SCN2A, SCN3A* and *SCN8A*

Na_V_1.1 represents one of several voltage-gated sodium channel subunits in which nearly all residues are structurally and functionally identical. As such, variants in one sodium channel can be analyzed by considering known effects in variants across the voltage-gated sodium channel gene family.^24^ Accordingly, we examined the literature for paralogous variants in *SCN2A, SCN3A*, and *SCN8A* and compared functional and phenotypic profiles.

Seven individuals were reported with disease-causing variants at paralogous sites, including *SCN8A* p.R1617Q (n=5), *SCN2A* p.R1626Q (n=1), and *SCN3A* (n=1).^25-31^ The individual with a *de novo SCN2A* p.R1626Q variant had a DEE presentation. The individual with a *de novo SCN3A* p.R1621Q had polymicrogyria and severe speech delay without epilepsy.^31^ The five individuals with the recurrent *SCN8A* p.R1617Q variant had DEE with seizure onset in the first few months of life (Table S1). Information on development was available for 3/5 individual and ranged between severe and profound intellectual disability.

For *SCN3A* p.R1621Q and *SCN8A* p.R1617Q, functional studies had been performed previously.^31-33^ Both variants showed slowing of fast inactivation, increased persistent current, and hyperpolarizing shift of voltage dependence of activation, leading to an overall gain-of-function (Figure 4). In neuron cell lines, the *SCN8A* variant was further found to have acceleration of recovery of fast inactivation, increased slope of fast inactivation, and increased resurgent current.^32^ In the non-epilepsy-related voltage-gated sodium channels *SCN4A* and *SCN5A*, disease-causing variants at the same codon also display similar overall gain-of-function functional effects.^34,35^ Thus, the electrophysiological footprint of these variants across sodium channel genes was nearly identical.

**Figure 4.**
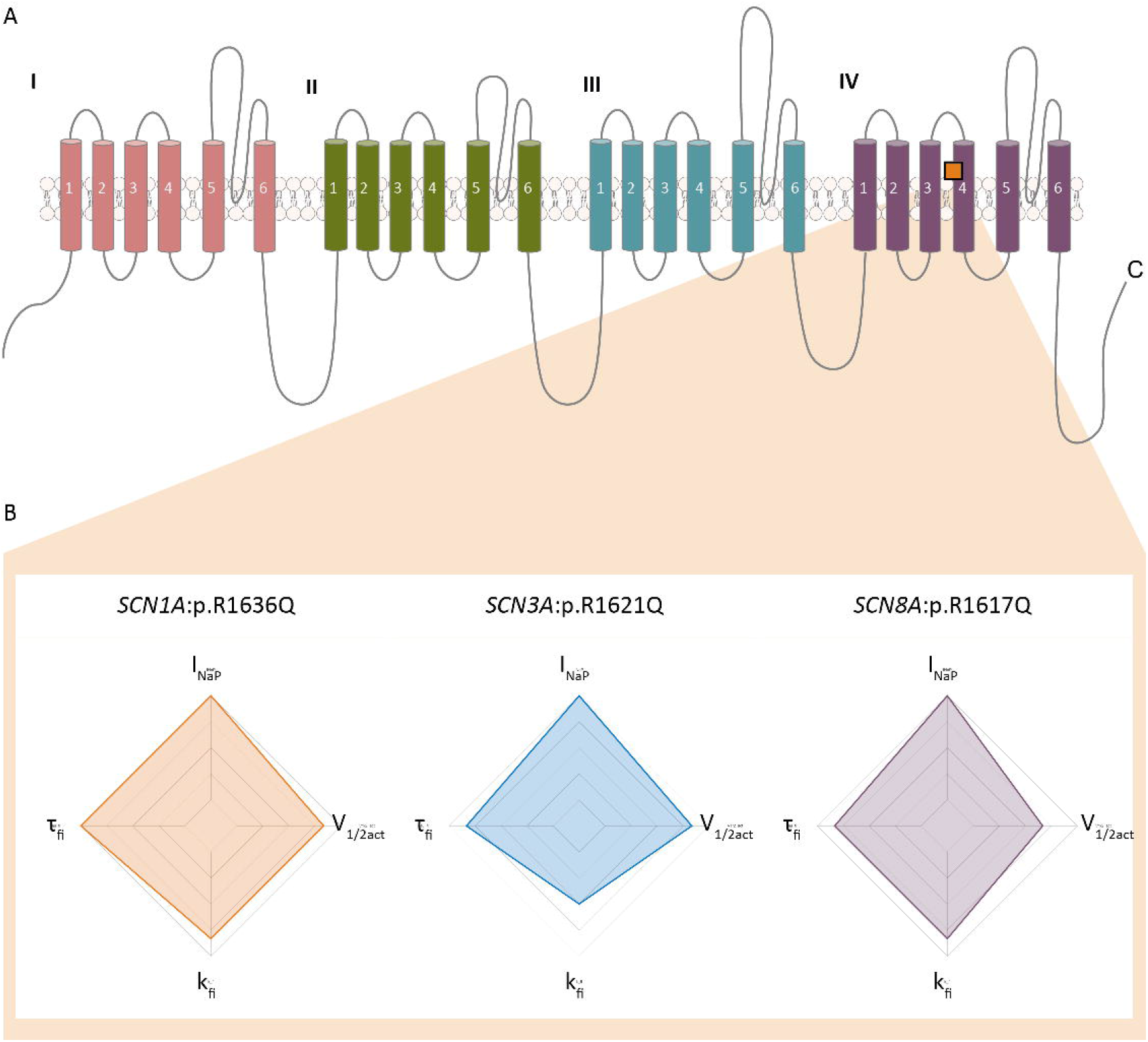
Functional features in *SCN1A* p.R1636Q and the identical variants in paralogous voltage-gated sodium channel genes. **(A)** Structure of the canonical voltage-gated sodium channel sequence showing location of the mutated arginine residue. **(B)** Functional deviations from wildtype scaled by severity. All variants show increased persistent current, slope factor of fast inactivation, time constant of fast inactivation, and voltage dependence of activation.

### Age of seizure onset for *SCN1A*:p.R1636Q is amongst the earliest in *SCN1A*-related disorders

Given the ultra-early-onset epilepsy phenotype in individuals with the recurrent p.R1636Q variant, we tried to understand how age of seizure onset compares to other individuals seen in our healthcare network. Also, we aimed to identify whether early clinical features might suggest the different functional consequence of this variant.

First, we retrieved information on age-of-onset in the study database for our Epilepsy Genetics Research Protocol (EGRP, Figure 5A). We identified 52 individuals with *SCN1A*-related disorders amongst the 2,356 participants in our study. Median age of seizure onset was 4.8 months with range 1 day to 4 years. Individuals with *SCN1A* p.R1636Q were below the second percentile for age of onset across all *SCN1A*-related disorders, representing the earliest ages of onset in our *SCN1A* study cohort.

**Figure 5.**
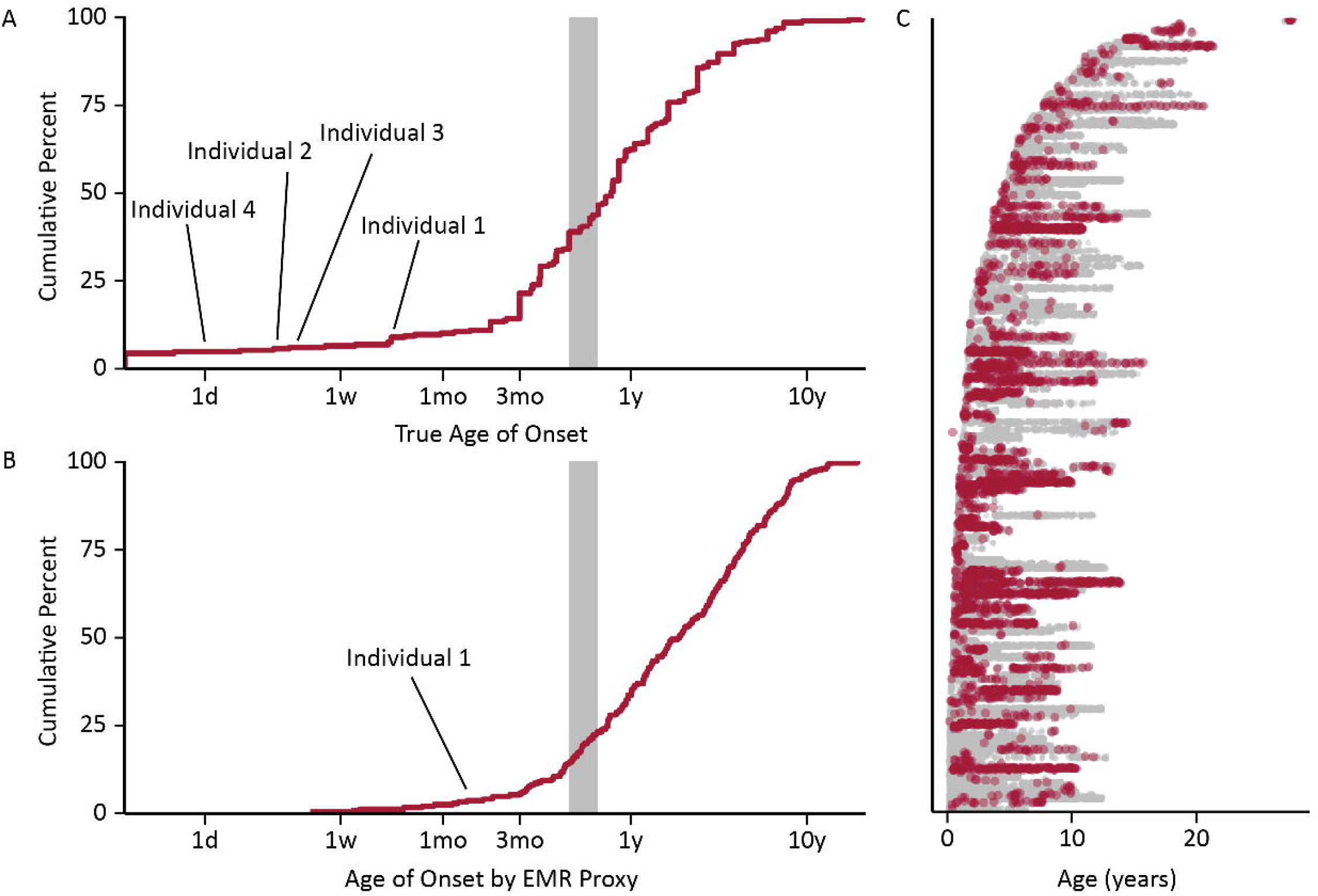
Electronic medical record analysis of individuals with *SCN1A*-related clinical notes. **(A)** 98,974 data-time points including 3,013 with explicit coding of a seizure diagnosis (red) across 1180 patient-years in the medical record. **(B)** True documented age of seizure onset in 52 individuals with *SCN1A*-related disorders. **(C)** Automatically extracted age of first seizure diagnosis in 213 individuals in the EMR showing that automated extraction yields a similar distribution to true onset and can identify ultra early-onset epilepsies related to *SCN1A* variants.

We then compared this true age of onset distribution to the ages of first recorded G40+ diagnosis, representing a proxy measure that can be automatically extracted from the electronic medical record. For all individuals with a mention of *SCN1A* and a G40+ diagnosis code, we further assessed when a G40+ diagnosis is first mentioned within a patient’s chart (**Figure 5B**). We identified 98,974 clinical notes across 213 individuals with epilepsy and *SCN1A* noted within the EMR, accounting for 1180 patient-years (**Figure 5C**). This automated capture recapitulated the identification of early-onset *SCN1A*-related epilepsy in routine care: while Individuals 2-4 initially presented at our healthcare system long after their first seizure, Individual 1 with the p.R1636Q variant had among the top 5% earliest appearances of G40+ diagnosis codes in the EMR (**Figure 5C**). This individual had an EMR age of onset of less than 1.3 months in contrast to the typical 6-9 months onset in Dravet Syndrome.

Of note, not all individuals with EMR records referring to *SCN1A* had a final diagnosis of an *SCN1A*-related epilepsy, but this search strategy allowed for an efficient way to obtain the references to a gene name within the entire healthcare network. We conclude from this analysis that EMR data can successfully identify individuals with ultra early-onset *SCN1A*-related epilepsies with putative gain-of-molecular-function variants.

## Discussion

Here, we report the clinical findings in four individuals carrying the recurrent SCN1A p.R1636Q variant and assess the functional consequences of this variant in heterologous systems. We demonstrate that individuals have much earlier seizure onset than the typical disease course in Dravet syndrome. Patient outcomes were variable with respect to seizure freedom and developmental outcomes, and the clinical response to sodium channel blockers was tentatively positive in those individuals where it was initiated early in disease course. As a functional consequence, we identify an overall gain-of-function due to hyperpolarized voltage dependence of activation and an increased persistent current.

We identify two major findings. First, we identify a clinical presentation in individuals with the recurrent p.R1636Q variant that is different from the previously reported phenotype. All individuals had early-onset epilepsy starting before the age of 2 months. We identified eight previously reported individuals with the recurrent p.R1636Q variant.^17,20-23^ In contrast to the ubiquitous presence of severe intellectual disability and hyperkinetic movement disorder in prior reports, the range of phenotypes in our patients was broader. We attribute these differences to the fact that our patients were recruited from ongoing care rather than retrospectively, in parallel to expansion of phenotypic spectra seen in many other genetic epilepsies such as SCN2A-, STXBP1-, and SCN8A-related epilepsies.^36-38^

Second, we show that the p.R1636Q variant results in an overall gain-of-function. Our electrophysiological analyses revealed a hyperpolarizing shift of activation and increased persistent current with a mild hyperpolarizing shift of inactivation. Though this reflects a mixed effect, the overall consequence is expected to be gain-of-function, in line with available functional and clinical information for identical disease-causing variants in the other epilepsy-related sodium channels, SCN2A, SCN3A, and SCN8A. In addition, paralogous variants have been shown to result in similar gains of function in non-neuronal sodium channel genes. Specifically, SCN5A p.R1623Q is associated with long-QT syndrome and the mutant channel exhibited gain of function due to significantly delayed inactivation properties while single channel analysis revealed that SCN5A R1623Q channels have significantly prolonged open times with bursting behavior.^39^ Moreover, a variant affecting the same codon in *SCN4A*, p.R1448C, was identified in an individual with paramyotonia congenita and displayed identical features to *SCN1A* p.R1636Q, namely hyperpolarization of both channel availability curves and prominent persistent sodium current. Notably, in parallel with oxcarbazepine in Na_V_1.1, ranolazine was shown to be effective for the Na_V_1.4 variant, reflecting its potency against persistent current.^40^

While most results have been performed in heterologous systems without the ability to assess the impact of neuronal firing, the relationship between phenotypes and biophysical abnormalities in *SCN1A* gain-of-function variants requires further examination. This is relevant given the different treatment strategies suggested by the various gain-of-function mechanisms. For example, oxcarbazepine stabilizes the fast-inactivated state of the Nav1.1 channel, while lacosamide acts on the slow inactivated state.^34,35^ Accordingly, for individuals with disease-causing gain-of-function variants in *SCN1A* demonstrating increased persistent current such as the recurrent p.R1636Q variant, lacosamide may potentially represent a more appropriate therapeutic choice explained by the specific pharmacodynamics of this medication. However, given the complex functional consequences of the p.R1636Q variant, neither sodium channel blocker will likely achieve a complete correction of the biophysical abnormalities.

In summary, we report clinical and electrophysiological features associated with the recurrent *SCN1A* p.R1636Q variant in four individuals at a single center. Adding to eight previously reported individuals, the recurrent p.R1636Q variant may represent the most common gain-of-function variants in *SCN1A*, possibly present in individuals previously considered to have atypical Dravet Syndrome as in some of the individuals in our study. Given the possibility that the identified gain-of-function effect can be approached at least partially with common sodium channel blocking anti-seizure medications, awareness of the “Philadelphia variant” and other gain-of-function variants is critical for targeted treatment in *SCN1A-*related disorders.

## Acknowledgements

This work was supported by The Hartwell Foundation Individual Biomedical Research Award and NIH NINDS K02 NS112600 to I.H., NIH NINDS U54 NS108874 (PI, Alfred L. George), and NIH NINDS R01 NS110869 and the Burroughs Wellcome Fund Career Award for Medical Scientists to E.M.G.

## Data Availability Statement

The data that support the findings of this study are available on request from the corresponding author. The data are not publicly available due to privacy or ethical restrictions.

## IRB Statement

This study was completed per protocol with local approval by the Children’s Hospital of Philadelphia (CHOP) Institutional Review Board (IRB 15-12226).

## Ethical Publication Statement

We confirm that we have read the Journal’s position on issues involved in ethical publication and affirm that this report is consistent with those guidelines.

