## Supplementary material for "The *SCN1A* Philadelphia variant – a gain-of-function mutation causing an early-onset epileptic encephalopathy": Table S1

|  | Individual 1 | Individual 2 | Individual 3 | Individual 4 | Individual 5 | Individual 6 | Individual 7 |
| --- | --- | --- | --- | --- | --- | --- | --- |
| Source | Willig et al  2015 | Zaman et al 2020 | Rauch et al 2012 | Ohba et al  2014 | Dyment et al 2015 | Kong et al 2015 | Larsen et al 2015 |
| Age (sex) | 2m* (N/A) | 15y (M) | N/A (F) | 2y (F) | 4y (F) | 3y (M) | 3y* (F) |
| Gene | *SCN2A* | *SCN3A* | *SCN8A* | *SCN8A* | *SCN8A* | *SCN8A* | *SCN8A* |
| Variant | c.4877G>A, p.R1626Q,  *de novo* | c.4861C>G, p.R1621Q,  not maternally inherited | c.4850G>A, p.R1617Q,  unknown | c.4850G>A, p.R1617Q,  *de novo* | c.4850G>A, p.R1617Q,  *de novo* | c.4850G>A, p.R1617Q,  *de novo* | c.4850G>A, p.R1617Q,  *de novo* |
| Seizure types (onset of first seizure) | N/A | T (4m), FIAS | N/A | FS (3m), GTC (6m) | M, AtA, GTC | FS, M, GTC | GTC (5.5m), C, T, AtA, M |
| EEG | N/A | Bilateral SW | N/A | Normal | Anterior mid-line frontal onset localization | Unilateral SW | Bilateral delta activity, unilateral SW |
| Brain MRI | N/A | Bilateral PMG | N/A | Normal | Normal | Normal | Normal |
| Neurological/physical exam | Arthrogryposis, pulmonary hypoplasia | Pseudobulbar palsy; severe ID | N/A | Profound global developmental delays; sitting and crawling | Developmental delay | Severe ID | Dystonia, hypotonia, NA |

**Table S1.** Clinical features in previously reported individuals with variants identical to *SCN1A*:p.R1636Q in other brain-expressed voltage-gated sodium channels.

At A: atypical absence, FIAS: focal impaired awareness seizure, FS: febrile seizure, GTC: generalized tonic-clonic, M: myoclonic seizure, NA: non-ambulatory, PMG: polymicrogyria, SW: spike wave, T: tonic *deceased
